# Aberrant regulation of RNA splicing in sunflower hybrids may underlie intrinsic incompatibilities

**DOI:** 10.1101/2020.09.08.287169

**Authors:** Chris C R Smith, Loren H Rieseberg, Brent S Hulke, Nolan C Kane

## Abstract

Alternative spicing is an integral part of gene expression in multicellular organisms that allows for diverse mRNA transcripts and proteins to be produced from a single gene. However, most existing analyses have focused on macro-evolution, with only limited research on splice site evolution over shorter term, micro-evolutionary time scales. Here we examine splicing evolution that has occurred during domestication and observe 45 novel splice forms with strongly transgressive isoform compositions, representing 0.24% of analyzed transcripts. We identify loci associated with variation in the levels of these splice forms, finding that many novel transcripts were regulated by multiple alleles with non-additive interactions. A subset of these interactions involved the expression of individual spliceosome components. These overdominant and epistatic interactions often resulted in alteration in the protein-coding regions of the transcripts, resulting in frameshifts and truncations. By associating the splice variation in these genes with size and growth rate measurements, we found that none of the individual splice variants affected these plant traits significantly, but the cumulative expression of all aberrant transcripts did show a significant reduction in growth rate associated with higher proportions of disrupted transcripts. This demonstrates the importance of co-evolution of the different spliceosomal components and their regulators and suggests that these genes may contribute to evolution of reproductive isolation as Bateson-Dobzhansky-Muller incompatibility loci.

**Author summary:** In multicellular organisms, it is common that segments of pre-mRNA molecules are physically removed, and the remaining segments are spliced back together. Through splicing alternative combinations of segments together, organisms produce various mRNA molecules, and thus multiple proteins, using the information encoded in a single gene. Here, we investigated the RNA of two sunflower genotypes, one wild and one domesticated, as well as the hybrid offspring resulting from a cross between the two genotypes. We found certain mRNA molecules that were spliced exclusively in the hybrids and were absent in the examined parental lines. These unique hybrid mRNAs were predicted to be consequential for the hybrids’ health, and thus represented a malfunction in the mechanisms that regulate splicing. These results improve our understanding of the genetic regulation of alternative splicing and how alternative splice forms evolve. Our findings may lead to further inquiries about how aberrant splicing promotes the formation of new species in nature.

## Introduction

Gene expression in multicellular organisms involves alternative splicing, which uses the information encoded in a single gene to create an assortment of mRNA transcripts. For example, as many as 150,000 unique transcripts were found in *Arabidopsis thaliana* (Huang et al., 2019a), a plant with only 27,466 protein coding genes (Cheng et al., 2017). Furthermore, relatively few splicing isoforms are conserved among plant species: only 14.7% of isoforms are conserved between *Arabidopsis* and rice, and only 16.4% are conserved between rice and maize (Severing et al., 2009; Zhang et al., 2015). Despite the large diversity of isoforms found in nature, our knowledge about the evolution of splicing, and the origins of novel splice types, is limited.

Recent studies have shown that RNA splicing may change over relatively short evolutionary timescales, e.g. 1,000-10,000 years (Thatcher et al., 2014; Zhang and Xiao 2018; Smith et al., 2018; Khokhar et al., 2019; Huang et al., 2019b; Lin et al., 2020). In domesticated sunflowers, divergent splice forms were identified that seemed to have arisen from standing variation (Smith et al., 2018). In this case, the divergent splice forms found in domesticated genotypes were also sometimes found in wild populations, suggesting that this variation existed at some frequency in the ancestral population before drift or selection brought certain splice forms to fixation in domesticated cultivars. Several questions remain: In hybrids, how often is splicing transgressive, leading to variation not seen in parental lineages? Are novel transcripts costly in hybrids? At a genetic level, what regulation appears to be associated with novel splicing?

Hybridization is one possible source of phenotypic variation in this case (Lewontin and Birch, 1966; Scascitelli et al., 2010). In regard to RNA splicing, a hybrid individual will inherit regulatory alleles from each parent and thus could create a set of transcripts that resembles that of either or both parents. On the other hand, it is possible that new alleles that were benign in one parent genetic background may interact with unfamiliar alleles in the opposite parent genome (Bateson, 1909; Dobzhansky, 1936; Muller, 1942), causing mis-regulation of splicing and the formation of novel RNA transcripts (Fig. 1).

**Figure 1.**
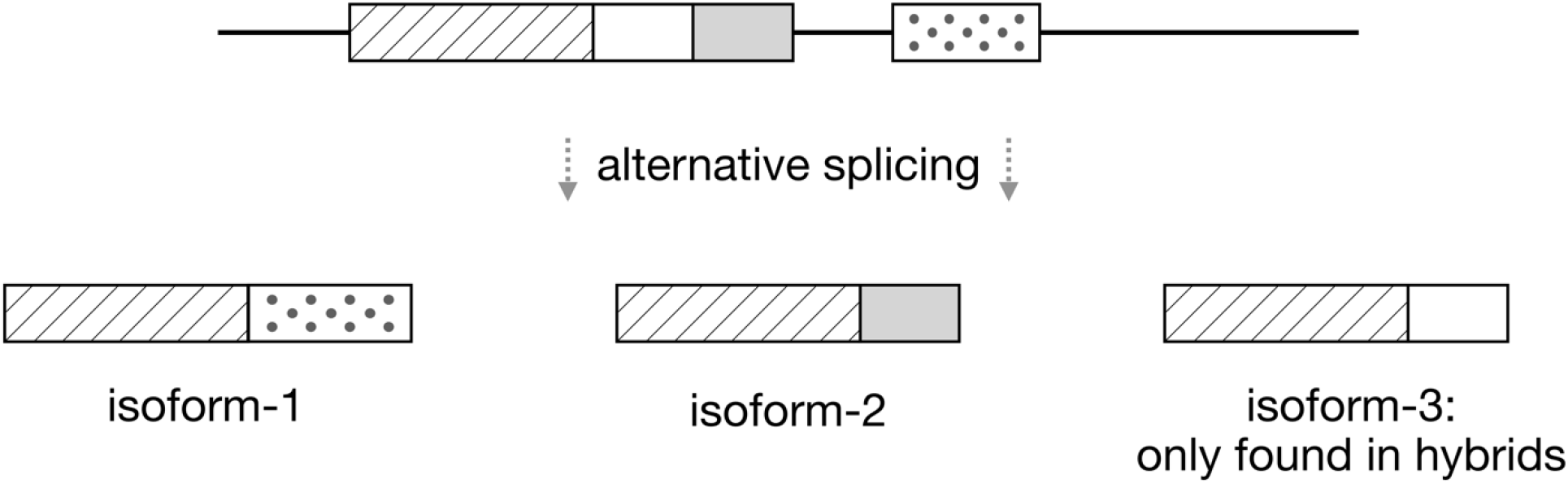
Alternative splicing diagram. The black line shows a DNA strand, and shaded boxes represent exon or retained intron segments. The bottom boxes are mature mRNA transcripts resulting from alternative splicing. We refer to theoretical isoform-3 as a novel, transgressive isoform because it is found exclusively in hybrids. Other isoforms could be transgressive if spliced at extreme levels relative to the parent taxa.

This phenomenon may be thought of as an example of transgressive segregation, because the hybrid phenotype is more extreme than- or in this case, nonexistent in-either parent (Slatkin and Lande, 1994). Transgressive segregation is not uncommon for a wide range of phenotypes, with 36-44% of examined traits in hybridizing systems showing some degree of transgression (Rieseberg et al., 1999; Stelkens and Seehauson, 2009). Transgressive segregation has also been observed in splice isoform variation (Scascitelli et al., 2010), although these observations have been limited to relatively few isoforms and the genetic regulatory mechanisms have not been identified. In general, the predominant genetic basis for transgression is complementary gene action, meaning that some proportion of hybrids may have alleles from each parent that combine additively to produce the extreme phenotype (see the first table from Rieseberg et al., 1999). Other causes of transgressive segregation include interactions between alleles at different loci, i.e. epistasis, or within a locus, i.e. overdominance.

Here, we investigated transgressive splice forms in the common sunflower, *Helianthus annuus*. We surveyed the entire transcriptome of one hundred recombinant inbred lines (RILs) derived from a wild x domesticated cross and mapped the genetic basis of transgressive splicing. Our findings help us understand how alternative splice forms evolve and how genetic incompatibilities contribute to the formation of new species.

## Results

### Characterization of novel splice forms in hybrids

We identified 45 isoforms that were spliced in at least one hybrid at a substantial level (≥ 1 transcripts per million; TPM), were absent in all six parent samples (zero reads), and appeared to each be an alternative splice form of a corresponding parent isoform (Table 1). We refer to these as “novel” isoforms to describe that they were not detected in the parent samples. These 45 isoforms represented 0.24% of analyzed transcripts after initial filtering. All exons and retained intron sequences in the novel transcripts were present in the Ha412-HO genome; this check supports that these transcripts represent alternative splice forms instead of sequence mutations, e.g. indels. Additional cases of transgressive splicing potentially exist, but these represented the most convincing examples in the current dataset.

**Table 1.** Trangressive isoform information. The fifth column lists all splice types between the novel isoform and parent isoforms: (IR), exon skipping (ES), or alternate splice site (AS). The open reading frame column lists truncated (T), elongated (E), otherwise different (D), or same/unaffected (S). The genetic basis column lists epistasis (E), overdominance (O), interaction with a spliceosome component that was differently expressed in the parents (S), marginally significant epistasis (e, by itself), single-QTL (q), and weak interactions between single-QTLs (qe).

| ID | Chrom., cM | Length (bp) | Annotation | Splice type | OR F | Genetic basis | Spliceosome component associations |
| --- | --- | --- | --- | --- | --- | --- | --- |
| 1 | 12,63.99 | 632 | cytochrome B561-1 | IR,ES,AS | E |  |  |
| 2 | 3,24.64 | 766 | sucrose phosphate synthase 2F | IR,ES,AS | T | E | RNA helicase DDX23/PRP28 |
| 3 | 16,42.46 | 1,231 |  | IR,ES,AS | T | O |  |
| 4 | 17,47.71 | 2,773 | ATPase E1-E2 type family protein / haloacid dehalogenase-like hydrolase | IR,ES,AS | E |  |  |
| 5 | 17,47.71 | 2,868 | ATPase E1-E2 type family protein / haloacid dehalogenase-like hydrolase | IR,ES | S | E |  |
| 6 | 6,29.29 | 1,638 | glycerol-3-phosphate acyltransferase | IR,ES,AS | S | E | pre-mRNA-splicing factor RBM22/SLT11, U6 snRNA-associated Sm-like protein LSm6/LSm7 |
| 7 | 17,69.49 | 1,251 | UBX domain-containing protein | IR,ES | T |  |  |
| 8 | 5,51.16 | 1,771 | FAD/NAD(P)-binding oxidoreductase | IR,ES,AS | T | E |  |
| 9 | 12,64.01 | 3,162 | ABC-2 type transporter | IR,ES | T | E | ATP-dependent RNA helicase DDX23/PRP28 1, nuclear cap-binding protein, pre-mRNA-splicing helicase BRR2, U5 small nuclear ribonucleoprotein helicase |
| 10 | 12,64.01 | 3,274 | ABC-2 type transporter | IR,ES,AS | T | E | U6 snRNA-associated Sm-like protein LSm6 |
| 11 | 8,67.05 | 2,701 | Mitochondrial transcription termination factor | IR,ES,AS | E | O | U4/U6 small nuclear ribonucleoprotein PRP31, small nuclear ribonucleoprotein G |
| 12 | 4,57.32 | 2,441 | Insulinase (Peptidase family M16) | IR,ES,AS | T | O | small nuclear ribonucleoprotein D2 |
| 13 | 7,10.48 | 672 | indeterminate(ID)-domain 2 | IR,ES | T |  |  |
| 14 | 16,50.99 | 633 | vascular plant one zinc finger protein | IR,ES,AS | S | (qe) |  |
| 15 | 14,25.78 | 1,259 | NAD(P)H dehydrogenase (quinone)s | IR,ES,AS | T |  |  |
| 16 | 15,57.81 | 1,309 | P-loop containing nucleoside triphosphate hydrolase | IR | T | E | U4 |
| 17 | 14,2.821 | 4,783 | P-glycoprotein 13 | IR | T | E |  |
| 18 | 4,50.94 | 1,197 | Leucine-rich repeat protein kinase | IR,AS | T |  |  |
| 19 | 11,59.05 | 604 |  | IR,ES, | E | EOS | PRP38 family protein |
| 20 | 11,44.40 | 1,536 | Aldolase superfamily protein | IR,ES,AS | E | E | PRP38 family protein |
| 21 | 2,80.65 | 368 |  | IR,ES | S | (q) |  |
| 22 | 7,11.01 | 1,557 | S-locus lectin protein kinase | IR,ES | S |  |  |
| 23 | 9,36.52 | 1,379 | RNI-like superfamily protein | ES,AS | T | (q) |  |
| 24 | 5,56.84 | 2,338 | ARM repeat superfamily protein | IR | T | O |  |
| 25 | 2,32.95 | 1,641 | Ikzf5 (DUF668) | IR,ES | T | (qe) |  |
| 26 | 3,29.63 | 2,660 | glycosyl hydrolase 9A1 | IR,ES,AS | E |  |  |
| 27 | 9,24.78 | 4,200 | FAM91 carboxy-terminus protein | IR,ES,AS | T |  |  |
| 28 | 12,50.13 | 1,356 |  | IR,ES,AS | S |  |  |
| 29 | 15,27.82 | 1,298 | Pectin lyase-like superfamily protein | IR,other | S | (q) |  |
| 30 | 8,67.22 | 827 |  | ES,AS | S | E |  |
| 31 | 11,47.26 | 1,429 | UDP-N-acetylglucosamine (UAA) transporter family | IR,ES | T | O | PRP38 family protein |
| 32 | 14,9.072 | 720 | Uncharacterized protein family (UPF0016) | IR | T |  |  |
| 33 | 4,50.55 | 1,166 | auxin F-box protein 5 | IR | T | OS | U1 snRNA |
| 34 | 16,91.09 | 2,858 | oligopeptide transporter 7 | IR,ES,AS | S | (e) |  |
| 35 | 17,26.11 | 1,297 | FkbM family methyltransferase | IR,ES,AS | T |  |  |
| 36 | 17,26.11 | 1,246 | FkbM family methyltransferase | IR,ES,AS | T |  |  |
| 37 | 12,63.75 | 1,187 | alanine:glyoxylate aminotransferase | IR,ES,AS | S |  |  |
| 38 | 12,63.75 | 1,660 | alanine:glyoxylate aminotransferase | IR,ES,AS | S | O | pre-mRNA-splicing factor SYF2, pre-mRNA-processing factor 40 |
| 39 | 12,63.75 | 1,259 | alanine:glyoxylate aminotransferase | IR,ES,AS | T | EOS | U1 snRNA |
| 40 | 9,24.77 | 2,213 | Inositol monophosphatase family protein | IR,ES,AS | E | EO |  |
| 41 | 16,52.35 | 1,431 | Ribosomal protein S18 | IR,ES,AS | S | E | peptidyl-prolyl isomerase H (cyclophilin H) |
| 42 | 15,37.53 | 1,822 | membrane protein | IR,ES,AS | S | E |  |
| 43 | 10,37.08 | 2,848 | Terpenoid cyclases/Protein prenyltransferases | IR,ES,AS | S |  | U1 snRNA |
| 44 | 10,39.00 | 2,271 | sister-chromatid cohesion protein 3 | IR,ES,AS | E | EO | pre-mRNA-splicing factor ATP-dependent RNA helicase DHX15/PRP43 |
| 45 | 7,9.847 | 2,173 | D-aminoacid aminotransferase-like PLP-dependent enzymes | IR,ES | D | (e) |  |

The length of novel transcripts ranged from 368 to 4,783 bp (median = 1,431 bp), which was longer than the most similar parent isoform in 96% of cases. The intron and exon sequences that differed between each transgressive isoform and the corresponding parent isoform ranged from 20 to 1,741 bp (median = 105 bp). The number of RILs with ≥ 1 TPM of each transgressive isoform ranged from 1 to 60 (median = 3). The proportion of alternative splicing composition at the gene-level was usually low for each novel isoform, although five were spliced at higher proportions (> 60%). In other words, the transgressive isoform was usually a minor isoform while the parent isoform remained the dominant transcript, but sometimes the novel isoform was spliced as the dominant isoform or as the only isoform. Some hybrids were more transgressive than others: all examined hybrids had some splicing of at least five different transgressive isoforms, but some had up to 18 different transgressive isoforms.

The transgressive genes had 1 to 11 isoforms in the parent samples (median = 4), suggesting that genes with more complicated splicing might produce novel isoforms more frequently. The splicing type that produced the novel transcripts - intron retention, exon skipping, etc.-was complex: because the majority of transgressive genes were alternatively spliced in the parents, the method of splicing that produced the novel transcripts must be characterized in relation to multiple alternative splice forms. Furthermore, comparisons between a transgressive isoform and an individual parent isoform usually involved more than one type of splicing. For example, intron retention and exon skipping sometimes occurred at different parts of the transcript simultaneously. To summarize, intron retention in the transgressive isoforms was the most common splicing type: 46% of comparisons between transgressive isoforms and corresponding parent isoforms involved this splice type. Other splice types included intron retention in the parent isoform (31%; percentages not exclusive), exon skipping in the parent isoform (42%), exon skipping in the transgressive isoform (37%), alternative splice site causing longer transgressive transcript (21%), and alternative splice site causing longer parent transcript (20%). Splice types are listed in Table 1.

Genes with transgressive splicing patterns were distributed across all 17 chromosomes. However, the number of transgressive isoforms spliced per gene was sometimes greater than one: three genes each had two novel isoforms, and a single gene had three novel isoforms. The majority of transgressive genes had similar splicing patterns between wild and domesticated parents, although 25% were differentiated between parent taxa. In Smith et al. (2018), splicing had diverged in 40% of analyzed genes. This indicates that divergence in splicing of a particular gene is not a prerequisite for the production of novel isoforms.

### Functional annotation of novel transcripts

Forty transgressive isoforms were homologous with 35 different *Arabidopsis* proteins with different functional annotations (Table 1). No two transgressive genes had the same *Arabidopsis* homolog; however, for each gene with multiple transgressive isoforms, the isoforms aligned to different splice forms of the same protein in a cleanly delineated manner. The annotations spanned a diverse array of functions, and no functional category appeared predominant. There were five novel isoforms that did not align well to any *Arabidopsis* proteins, two of which had a parent isoform with a homolog. The annotations for these were translation initiation factor 3 subunit I and hypothetical protein.

Most transgressive isoforms aligned to the same protein splice variant as one of the corresponding parent isoforms. This result is consistent with the transgressive isoforms being previously uncharacterized, although it is reasonable to suspect that this pattern is due to limitations of the alignment algorithm, or a less than comprehensive protein database. Only two novel isoforms aligned to different proteins than the parent isoform. In one of these cases, the transgressive isoform aligned to a different protein splice variant than the parent isoform, but they had the same annotation: uncharacterized protein family UPF0016, a suspected calcium transporter (Demaegd et al., 2014). In the other case, the transgressive isoform aligned to a protein with a different accession than the parent isoform, but only a subtly different functional annotation. The parent isoform was annotated as an F-box/RNI-like superfamily protein, while the transgressive isoform was most similar to the RNI-like superfamily protein. The transgressive isoform homolog is annotated as being involved in the degradation of KRP1/ICK1, which is not a function of the parent variant.

### Open reading frame analysis

The longest open reading frame (ORF) had been altered in most novel transcripts. Specifically, 31 novel isoform ORFs were different versions of the corresponding parent ORF, or different than all parent ORFs if the parents spliced multiple isoforms. Twenty-two were shorter than the most similar parent ORF, eight were longer, and one novel ORF was the same length as the parent version but differed in a six amino acid sequence on the 5’ end. Twenty-five of the transgressive ORFs differed by > 25 amino acids compared to the most similar parent ORF. In 13 novel splicing cases, an intact parent ORF was present in the transgressive isoform; in these examples, we assumed that the overall function of the gene was unaffected. In the remaining transgressive gene, also the shortest transgressive gene identified, no ORF was identified in the parent isoform or the novel isoform.

### Novel splicing effects on stress response

We evaluated potential relationships between transgressive splicing and 15 measurements related to hybrid stress response. The measurements included: (i, ii) the number of days until wilting and the number of days until death after watering ceased; (iii, iv) soil water content at two timepoints; and (v - xv) a set of correlated measurements related to plant size including: height at three timepoints, dry and wet weight, longest leaf length and width at two time points, and most recent fully expanded leaf length and width. The transgressive isoform proportions were scaled and averaged to obtain a cumulative transgressive isoform value for each RIL. Three of the size-related traits had relationships with transgressive splicing: most recent fully expanded leaf width (*p* = 0.01; Fig. 2A), dry weight (*p* < 0.01; Fig. 2B), and wet weight (*p* = 0.02; *r*^2^ = 0.06). This analysis predicted the most transgressive seedlings to have 47% less mass (dry weight) compared to the least transgressive seedlings, although transgressiveness only partly explained variation in plant size.

**Figure 2.**
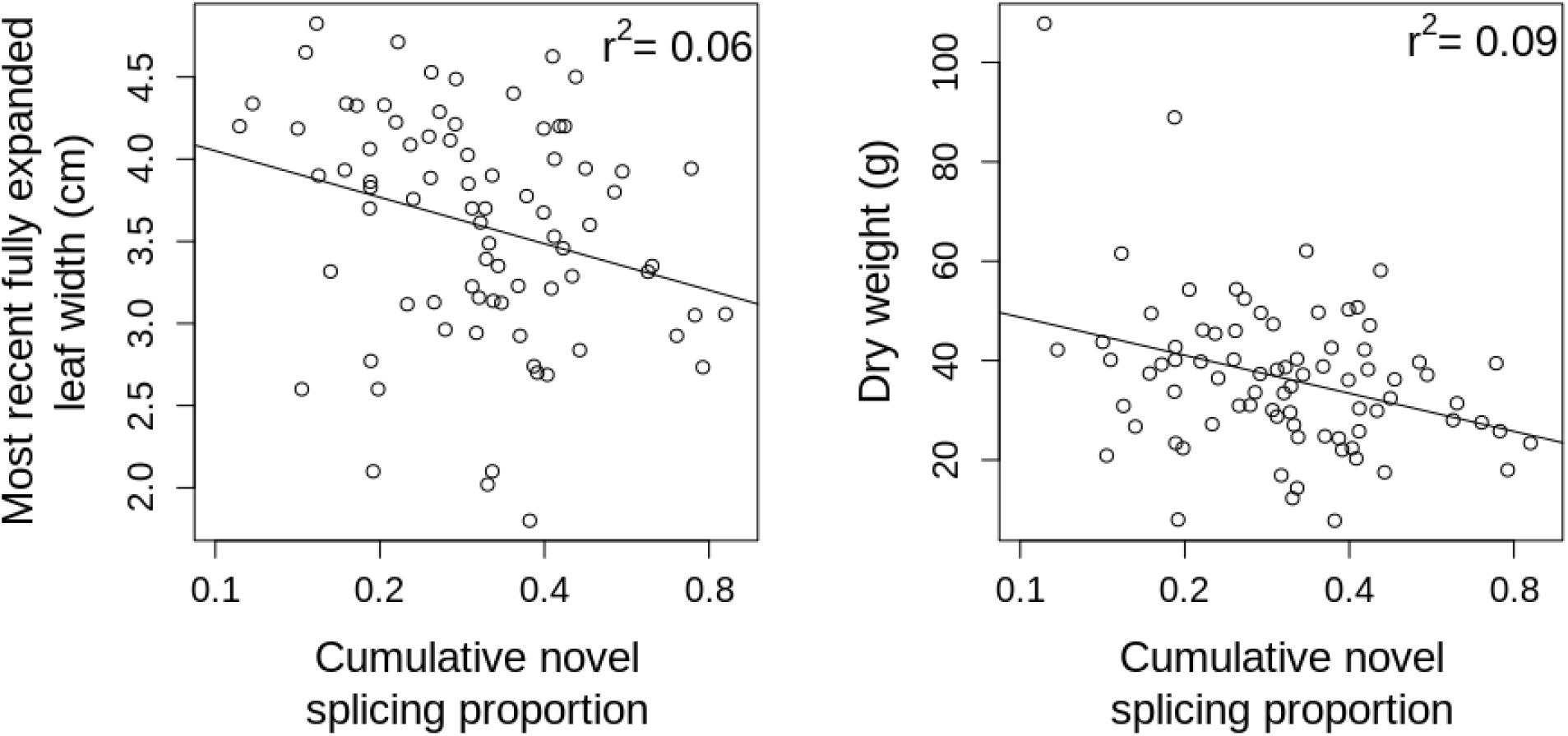
Stress response phenotypes versus transgressive splicing proportion. The x-axis is stretched to show log scale.

Additionally, we applied a principal component analysis to the eleven size traits, and the first principle component (88% of variance explained) was correlated with cumulative transgressive splicing (p = 0.01; *r*^2^ = 0.06). No individual isoform splicing levels had significant correlations with plant size or other measurements after correcting for multiple tests. Genome wide heterozygosity was not significantly correlated with plant size or other measurements.

### Overdominant regulatory loci were associated with novel splicing

To explore the genetic basis of each transgressive isoform, we scanned the genome for quantitative trait loci (QTLs) that had statistical associations with transgressive splicing. We detected QTLs for 19 different novel isoforms; these QTLs may regulate novel isoform splicing (Fig. 3, top row; Table 1). Most QTLs were associated with a single transgressive splice form, but seven QTLs each regulated two to five novel isoforms. The variance explained by each QTL ranged from 13% to 95% (median = 37%). The genomic position for most QTLs (85.7%) was more than 5cM away from the gene with transgressive splicing.

**Figure 3.**
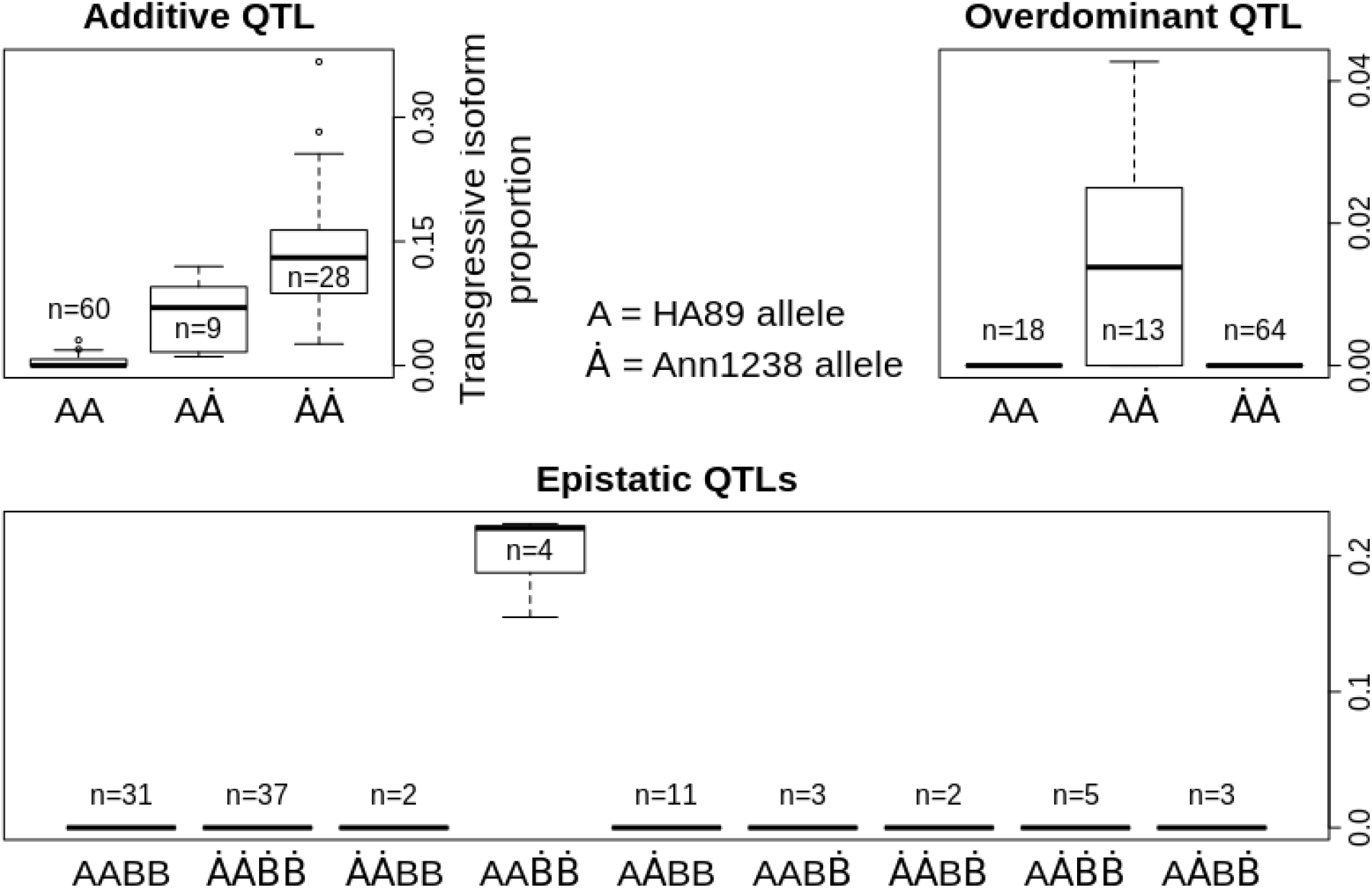
Transgressive splicing proportion for each genotype at example QTLs. (top left) Proportion of novel isoform #31 at an additive QTL (Chr. 11, 47.79 cM). (top right) Proportion of novel isoform #38 at an overdominant QTL (Chr. 13, 71.54 cM). (bottom) Proportion of novel isoform #2 for a pair of epistatic QTLs (Chr. 13, 70.78 cM; Chr. 16, 3.47 cM).

For each of the identified QTLs, we examined whether the heterozygous genotype was associated with transgressive splicing. Although the mapping population was inbred, the RILs retained heterozygosity at 10% of genomic sites on average. Eleven novel isoforms had QTLs with overdominance effects: individuals that were heterozygous at the QTL had a higher proportion of the transgressive isoform than either homozygous genotype group (Fig. 3, top right; Table 1). This indicates that alleles from each parent interacted non-additively to produce the transgressive splice type.

For only one novel splice form, the genotype at the QTL associated with transgressive splicing was homozygous Ha89. The homozygous Ann1238 genotype was associated with transgression for all remaining QTLs. Although fourteen novel isoforms were regulated by more than one QTL, in these instances each individual QTL-effect came from the same parent allele or overdominance. Therefore, the identified QTLs did not show evidence for complementary gene action, where alleles from both parents have effects that combine additively to produce the transgressive isoform.

### Epistasis underlies transgressive splicing

Next, we explored whether alleles at different loci interacted non-additively to produce the transgressive splice type. Irrespective of the single-QTL scan (above) we scanned the genome for pairs of loci where the effect of one locus depended on the genotype at the other locus. Sixteen novel splice forms could be explained by interacting regulatory loci (2.2 × 10^−20^ < p < 3.1 × 10^−11^; Fig. 3, bottom panel; Table 1). The number of epistatic QTLs per novel isoform ranged from 1 to 9 (median = 1.5). Most epistatic QTLs were *trans*, although 11.4% of interactions involved a QTL within 5cM of the gene with transgressive splicing.

It was revealing that significant epistasis was detected for seven novel isoforms with QTLs identified in the initial single-locus scan. Three novel isoforms had single-QTLs with genomic positions that overlapped with an epistatic QTL. This suggests that some of the single-QTLs may have been initially detected because of epistatic interactions with other loci. As a final check, we inspected whether the previously identified single-QTLs interacted with each other. Of the 14 novel isoforms that were each regulated by two or more single-QTLs, there were two pairs of interacting QTLs that affected the same novel isoform (1.4 × 10^−8^ < p < 3.3 × 10^−6^). Nine additional QTLs pairs affecting eight novel isoforms showed evidence of weak epistasis (1.1 × 10^−3^ < p < 3.8 × 10^−2^).

### Associations with particular spliceosome components

The spliceosome is a protein and snRNA complex that is responsible for the removal of introns and splicing-together of exons during alternative splicing. Using genomic resources for *Arabidopsis* to identify spliceosome genes in sunflower, we examined which individual spliceosome components were involved in transgressive splicing. We found associations between the expression level of individual spliceosome components and splicing of 16 different transgressive isoforms (Fig. 4; Table 1). While each novel isoform usually had only one spliceosome component association, five novel isoforms each had more than one association, and four spliceosome components were each used by more than one novel isoform. For example, the expression of U1 spliceosomal RNA was positively correlated with splicing of three transgressive splice forms. Our novel isoforms were not spliceosome RNA or protein components, themselves.

**Figure 4.**
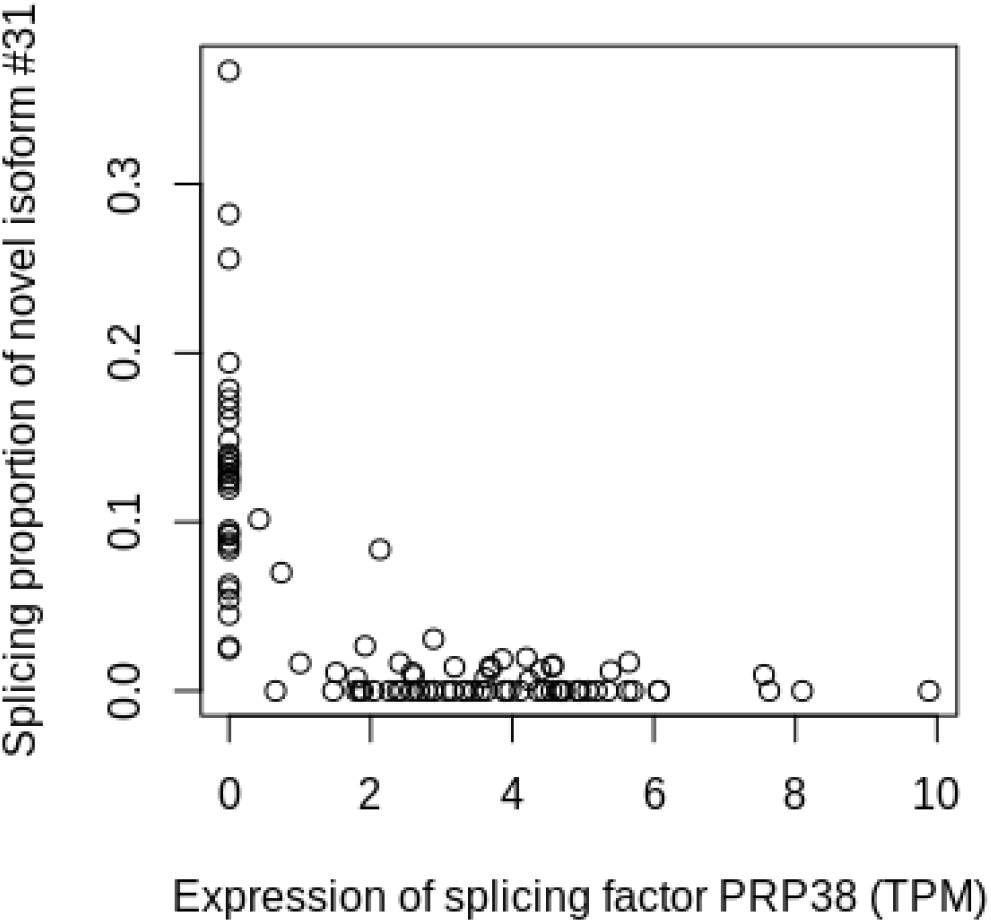
Novel isoform splicing versus spliceosome component expression.

We investigated which type of regulatory factor, QTLs or spliceosome component expression, was more important in the splicing of novel isoforms. There were 15 novel splicing examples where the transgressive isoform was associated with both (i) at least one identified regulatory QTL and (ii) at least one spliceosome component. Using a linear model with individual variables for each spliceosome component and QTL associated with a novel isoform, we partitioned the sum of squares explained by each factor. For six novel isoforms, the expression of the spliceosome components explained more variance in splicing than the QTLs. This pattern of variance propagation suggests that some-but not all-spliceosome components were more closely involved in novel isoform production. The QTLs in these six cases affected novel splicing less directly, perhaps through regulation of the spliceosome.

Next, we inspected whether the QTLs primarily regulated spliceosome components instead of directly regulating novel splicing itself. This was achieved by regressing each QTL against each spliceosome component, and against novel isoform splicing. There were eleven QTLs that explained more variance in the expression of a spliceosome component than the splicing of the novel isoform. These QTLs were associated with four different novel isoforms.

Last, we found statistical interactions between spliceosome components and QTLs regulating the same novel isoform in eight novel splicing cases (3.2 × 10^−35^ < *p* < 0.03), indicating that the two effects were sometimes interdependent. Looking back to the parental samples, two of the spliceosome components that interacted with QTLs had differentiated gene expression between parental genotypes (1.3 × 10^−3^ < *p* < 8.8 × 10^−3^). The latter signals that the parental genotypes have heritable variation in spliceosome expression that could lead to genetic incompatibilities in the hybrids.

## Discussion

### The evolution of new exons

Much of what we know about alternative splicing evolution describes changes that occurred over large evolutionary timescales, for example rates of intron gain and loss and differences in intron numbers among widely divergent taxa (Lynch and Walsh, 2007). The analysis presented here shows that RNA splice sites can evolve in a small number of generations through rewiring of existing regulatory networks. This contrasts with an existing model for splice site evolution predicting that mutations in the spliced gene sequence itself would be the main source of new exons (Keren et al., 2010). Another interesting pattern is that most of the affected genes had relatively high isoform diversity and complex splicing. This suggests that genes with higher splicing complexity may have a higher probability of creating new splice forms.

Four genes with novel splicing (novel isoforms 16, 23, 32, 33) each had a single mRNA form in the parental samples. These parental transcripts each aligned to the reference DNA in multiple pieces, indicating that they were spliced constitutively. These cases are of particular interest because they might tell us something about how alternative splicing evolved from constitutive splicing (assuming that genes with introns were an intermediate evolutionary state; Ast 2004). These four novel isoforms were usually spliced at low levels in the hybrids, but one isoform was the predominant splice type in the hybrid samples. For three of these genes, the new alternative splice form retained an intron, which is consistent with intron retention being the most common splice type in plants (Chamala et al., 2015). In the fourth gene, the alternative splice type was a combination of a newly incorporated exon and an alternative splice site. The latter case fits a model where a new exon originates and then may evolve to become more frequent. The gene sequence being spliced did not appear to be mutated in examined samples, which instead suggests a regulatory change. Indeed, we found QTLs associated with three of the above novel splice forms.

The longest ORF was inferred to be compromised in half of the novel isoforms in the current study, by frameshifts and truncations. We assume that frameshifts in mRNAs are costly for the organism, for example by affecting protein stability or binding domains, or by causing more serious disorders; although, nonsense-mediated decay may reduce some of these effects. On the other hand, new splice sites may occasionally lead to novel protein functionality, which could be beneficial if combined with the right adaptive landscape. Most of the novel isoforms in the current study had the same closest *Arabidopsis* homolog as the corresponding parental splice form. Therefore, the function of the novel isoforms may be conserved, or the function may be altered but as of yet uncharacterized. Experiments are needed to evaluate the function of the novel transcripts. As described by Keren et al. (2010), in cases where the ancestral isoform is still present, a non-coding mRNA or non-functional protein may have inherent regulatory function through limiting the production of functional proteins from the same gene.

A novel transcript might be thought of as qualitatively different than the parental transcript, especially if the corresponding protein function is altered. However, most novel transcripts in our dataset were spliced at low levels and the parental isoform remained the predominant form in the hybrids. Therefore, the presence of a new splice site may represent only a gradual step in the transition in alternative splicing for a gene. Simultaneously, recently evolved splice sites might endure a natural trial phase during which selection decides whether or not to favor the new splice type. Through this process, changes in splicing would evolve gradually, similar to other quantitative traits.

### The genetic basis of novel splicing

Novel transcripts in the current study were found in hybrid samples and absent in the parental samples, which is an extreme example of transgressive segregation. This type of inheritance indicates three possible genetic bases for the novel isoforms: epistasis, overdominance, or complementary gene action; although complementary gene action is generally accepted as the more common genetic basis for transgressive segregation (deVicente and Tanksley, 1993; Rieseberg et al., 1999; Stelkens and Seehausen, 2009). In the current study, QTLs were detected for 64% of novel isoforms. We did not experimentally verify any genetic associations.

Epistatic interactions explained 36% of novel splicing cases. The Rieseberg et al. (1999) review reported only one example of transgressive segregation caused by epistasis, however many of the reviewed studies did not test for epistasis. Stelkens and Seehausen (2009) hypothesized that epistasis and complementary gene action would explain most transgressive traits. This genetic basis for transgressive splicing is intuitive, because alternative splicing ordinarily relies on epistatic gene networks for the assembly and regulation of the spliceosome. In fact, we found that the expression levels of particular spliceosome proteins and spliceosome RNAs were correlated with the production of novel isoforms. In a subset of cases, the detected QTLs were more tightly associated with the expression of certain spliceosome components than the novel isoform itself, suggesting that the QTLs may regulate expression- or splicing- of the spliceosome. Furthermore, QTLs often had nonadditive interactions with spliceosome components, some of which had divergent gene expression in the parent taxa. Biologically, interactions involving the spliceosome may represent a specific type of hybrid incompatibility that leads to aberrant splicing.

In our dataset, inbreeding left relatively little heterozygosity in the RILs; nonetheless, we found that 24% of novel splice forms could be explained by overdominance at one or more QTLs. Overdominance, here, does not suggest that the hybrid phenotype is superior or better adapted. Few other transgressive phenotypes were explained by overdominance in the Rieseberg et al. (1999) review, but Stelkens and Seehausen (2009) reported many examples of heterosis in F1 hybrids. We would expect novel isoforms caused by overdominance to be even more common in natural hybrid zones due to higher heterozygosity. Transgressive phenotypes caused by overdominance are ephemeral, except in the case of allopolyploidy, because the heterozygous genotype cannot become fixed.

We did not find evidence for complementary gene action, where alleles from both parents have effects that add together to cause transgression. Although, it is possible that many undetected loci with small effects could contribute to transgressive splicing. Unexpectedly, individual QTLs were identified for six novel isoforms that could not be explained by overdominance or interactions with other loci. This result highlights a limitation in our ability to detect regulatory loci, because otherwise we would expect to see the aforementioned transgressive isoforms in one of the parental lines. It was illuminating that some of the QTLs detected in the single-locus scan had significant interactions with other regulatory loci; the remaining single-QTLs may have interacted with the genomic background more subtly, involving many small interaction effects. In fact, marginal epistasis was detected for three additional novel isoforms (p < 0.1 before correcting for multiple tests). Alternatively, a higher-order interaction scan might be required to see some kinds of epistasis. We did not find interactions or other QTLs involving cpDNA.

### Implications for speciation

Following a period of allopatry, mutations that were benign in one parent genetic background may be incompatible with the opposite parent genome. The field of speciation has produced a wealth of theory regarding this type of postzygotic incompatibility (Orr 1997; Orr and Turelli 2001; Turelli et al., 2001; Coyne and Orr 2004). Although BDM incompatibilities have maintained a central position in speciation theory, empirical evidence for this type of reproductive isolation has only begun to accumulate in recent decades (e.g. Brideau et al., 2006; Cattani and Presgraves, 2009; Anderson et al., 2010; Matute et al., 2010; Moyle et al., 2010). Empirical evidence for BDM incompatibilities is infrequent partly because sterile hybrid lineages are difficult to study and epistasis is difficult to detect. We still don’t fully understand how common BDM incompatibilities are, what types of genes are involved, and how important they are to allopatric speciation.

A breakdown in splicing regulation that leads to frameshifts in mRNA transcripts may be considered an incompatibility if it contributes to reproductive isolation. It is not uncommon for abnormal splicing of a single gene to be harmful: 10-15% of mutations underlying human diseases affect splicing (Nissim-Rafinia and Kerem, 2002). Although reproductive isolation was not directly tested in the current study, splicing malfunction led to mRNA frameshifts, and disrupted splicing was negatively correlated with seedling growth rate. We point out the feasibility of a scenario where an epistatic interaction involving the spliceosome simultaneously affects splicing of many loci and could lead to reduced hybrid fitness. Other links have been found between alternative splicing and adaptive radiation: Terai et al. (2003) showed that the putative pigmentation gene *hagoromo* has increased splicing diversity in cichlid species, in particular ones that have recently radiated, and different species exhibited fixed splicing differences. Also in cichlids, Singh et al. (2017) showed that alternative splicing patterns had diverged more dramatically between species than gene expression levels, and genes with differentiated splicing were associated with jaw specialization. Last, the noteworthy *Agouti* gene produced multiple isoforms in the skin of deer mice, and one particular isoform was associated with light-sand camouflage (Mallarino et al. 2017).

Future research is warranted to investigate splicing malfunctions in other hybridizing taxa, particularly between wild populations or species. Speciation researchers often test for epistasis associated with a particular reproductive isolation phenotype (e.g. sterility). If disrupted splicing is common in other taxa, researchers will have access to many potential targets for studying negative epistatic effects. Also, expression QTLs tend to have a high signal to noise ratio, which may improve detection of regulatory loci for splicing phenotypes. Most transgressive splicing cases in our study involved genes where splicing had not differentiated between parental lineages. This is consistent with the findings of Stelkens and Seehausen (2009), who hypothesized that stabilizing selection-not divergent selection-would favor transgressive phenotypes; however, that hypothesis was based on a model of complementary gene action, which we did not find evidence for. Contrasting predictions have been made about the relationship between genetic divergence and the frequency of transgression (Rieseberg et al., 1999; Stelkens and Seehausen., 2009). Additional research is necessary to determine what factors predict transgressive splicing.

## Materials and methods

### Plant material and RNA Sequencing

The raw sequence data from Smith et al. (2018) were reanalyzed in the current study. Ha89 is an inbred line commonly used in research and breeding (US Department of Agriculture Ames 3963). Ann1238 was derived from material collected at Cedar Point Biological Station, Keith County, Nebraska. Three seedlings from each parental accession and each of 100 sixth-generation recombinant inbred lines were grown in greenhouse conditions to address stochastic variation in expression. Above-ground tissue was frozen in liquid nitrogen, and total RNA was extracted following standard protocols. Nonnormalized Illumina RNA-sequencing (RNA-seq) libraries were sequenced on a HiSEq2000 system.

### Identification of transgressive splicing

We built a *de novo* transcriptome assembly including all parental samples and RILs using the program Trinity (v. 2.8.1) with default settings. This assembly differed from the Trinity assembly from Smith et al. (2018), because the previous assembly included only the parental samples. Henceforth, all data are presented for the first time. Custom scripts were used for all of the following steps and analyses, except when existing software is explicitly named. All custom code is available at https://github.com/chriscrsmith/SunflowerTransgressiveSplicing.

After completion of the assembly, the following initial steps were used to verify the assembled transcripts. All transcripts were aligned to the Ha412-HO genome (Staton and Rieseberg, 2019; Todesco et al., 2020), a domesticated sunflower genotype, using *BLAST* (McGinnis and Madden, 2004). Blast hits with percent identity below 95% were ignored. We required the entire transcript to have aligned ambiguously to the same genomic region, but with up to 10% of the transcript length missing from the ends. Additionally, the genomic alignment was required to be at least 50% single-copy-we delineated single copy sites by aligning whole genome sequencing data to the Ha412-HO reference and visualizing the distribution of read depths per site (S1 Text). If isoforms that were assigned the same gene-ID by Trinity aligned to different genomic locations, then we considered them as having been transcribed from different genes for all subsequent analyses.

Next, we used the following procedure to distinguish alternative splice forms assigned by Trinity from alternative alleles. We created a multiple sequence alignment of all isoforms from a gene using MUSCLE (v. 3.8.31), or, if only two isoforms, a pairwise sequence alignment using the EMBOSS (v. 6.6.0.0) *needle* program. For each pair of transcripts in the alignment: if at least one insertion/deletion or substitution larger than our minimum exon size cutoff-25 bp-was present, we considered the transcripts to be legitimate alternative splice forms. If any shorter runs of differences, e.g. a single nucleotide polymorphism (SNP), were present, we considered the transcripts to be alternative alleles of the same splice form; except, if both a > 25bp run and a shorter run were present, we deemed the relationship ambiguous.

We then applied the following filters to identify transgressive isoforms. For each potential transgressive isoform, we required: (i) all parents to have zero reads of the transgressive isoform and any alternative alleles, (ii) at least one hybrid to have substantial expression-> 1 TPM-of the transgressive isoform, and for there to be (iii) at least one alternative splice form with > 1 TPM in every parent. Because these isoforms are expressed at a substantial level, they are presumably distinct from stochastic splicing errors or noise.

### Testing for splicing differentiation

To test for divergent splicing composition between wild and domesticated sunflower genotypes, we first applied an isometric log ratio transformation to the proportions of each isoform (Smith et al., 2018). Then we used a t-test, or a MANOVA in the case of more than two parent isoforms, to test for a difference in splicing composition.

### Gene annotation

Transcripts were aligned to the Araport11 table from The *Arabidopsis* Information Resource (Berardini et al. 2015) using BLASTX (McGinnis and Madden, 2004) with e-value < 10^−20^ cutoff. The best hit was retained for each transcript.

### Open reading frames

The program ORFfinder (National Center for Biotechnology Institute website; www.ncbi.nlm.nih.gov/orffinder/) with default settings was used to identify open reading frames for each novel isoform and all parent isoforms corresponding to the same gene.

### Growth rate experiment

Ha89 × Ann1238 RIL seedlings, and inbred parental lines, were grown in a common environment in greenhouses at the University of British Columbia in the spring of 2011, with plants watered and fertilized daily. Plant height, leaf lengths and widths, and other measurements were taken at 4 weeks and 6 weeks after germination. At 6 weeks, watering was stopped entirely, with daily observations checking for wilting and death of each plant. Soil moisture content was measured with a HydroSense CS620 soil moisture system (Campbell Scientific) at 6 days after the initiation of the drought and on the day of first observed wilting for each plant. Plant measurements from this experiment are in Table S1.

### Single nucleotide variant identification and filtering

SNPs were identified by aligning RNA-seq reads to the Ha412-HO transcriptome (Renaut et al., 2013) by using bwa mem v0.7.15. SNPs were called by using SAMtools v1.4.1. We filtered SNPs that aligned to non-single-copy genomic regions (S1 Text) and marked genotypes with fewer than ten read depth as missing data. After the initial filters, we kept 66,893 SNPs that showed fixed differences between wild and domesticated parental samples. Next, we obtained genetic map positions for each fixed parent SNP by aligning transcriptome contigs to the Ha412-HO genome using BLAST (e value ≤10−20; ID ≥90%) and linearly interpolating cM positions from the Ha412-HO genetic map (Staton and Rieseberg, 2019; Todesco et al., 2020). Markers that aligned well were examined in the RILs, and genotypes with fewer than ten reads were marked as missing. Markers were excluded if present in fewer than 35% of RILs, resulting in 13,874 filtered SNPs. Last, we applied a conservative imputation step to the filtered SNPs: if a marker with missing data occurred between two markers with the same genotype that were within 10cM, we assigned the same genotype to the missing marker. Otherwise, missing genotypes were left as missing. This step filled in 31% of missing data, and 1.4% of genotypes overall.

### QTL mapping

To explore the genetic basis of each transgressive isoform, we used the program R/qtl (Broman et al., 2003) to conduct standard interval mapping with a non-parametric model. The mapped phenotype value was the proportion of transgressive isoform spliced relative to overall gene expression. The densest genetic map grid (*step* = 0) was used, and the genotyping error rate was set to zero (*error*.*prob* = 0) because the genotypes were already carefully filtered (above). We generated a null distribution of LOD scores for each novel isoform being mapping by permuting the phenotypic measurements relative to the genotypes: for 10,000 iterations, the splicing measurements were shuffled before applying the QTL scan, and the largest LOD score was recorded from each permutation; the 95^th^ percentile of this distribution was used as the significance threshold (Broman and Sen, 2009). We allowed a single QTL per chromosome and assigned the QTL peak as the marker position with the highest LOD score. We delineated the QTL region as the range of SNP markers with LOD scores within 25% of the peak. Overlapping QTL regions from different novel isoforms were consolidated for reporting the total number of QTLs.

For each QTL identified in the above scan, we calculated the degree of dominance at the QTL peak. The degree of dominance is the ratio of the dominance effect to the additive effect, the former being the difference between the mean phenotype of the heterozygote group and the midpoint between the mean phenotypes of each homozygote group, and the latter being half the difference between the mean phenotypes of the two homozygous groups (Kenney-Hunt, 2006; Ishikawa, 2009). A degree of dominance equal to zero means perfect codominance, 0 to 1.5 means one allele is dominant, and greater than 1.5 was interpreted as overdominance.

### Epistasis scan

For each novel isoform, we scanned the genome for pairs of loci where alleles from each parent were interacting to cause transgressive isoform splicing. The proportion of transgressive isoform spliced relative to overall gene expression was used as the phenotype value, and if an individual had insufficient expression (< 1 TPM) at the gene-level they were assigned missing data. We limited the search to pairs of loci on different chromosomes that met the following criteria: (i) at least three individuals were represented that had non-missing phenotype data and were homozygous for the Ha89 allele at both loci-notated *AABB*-, (ii) at least three individuals were represented that were homozygous for the Ann1238 allele at both loci- *ȦȦḂḂ*-, (iii) at least three individuals were represented for at least one mixed genotype - *ȦȦBB, AAḂḂ, AȦBB, AABḂ, ȦȦBḂ, AȦḂḂ*, or *AȦBḂ*. Encoding the genotypes as allele dosages-0, 1, or 2-, we tested for an interaction effect between markers using a multiple linear regression. We used a Bonferroni correction to obtain a significance threshold for the epistasis scan: the total number of tests for all 45 phenotypes was 1.61 × 10^9^, and the significance threshold became p < 3.11 × 10^−11^. Last, we applied the following post hoc filter to avoid outlier effects: if the most transgressive genotype group had fewer than three individuals represented, we counted an otherwise significant test as non-significant. We allowed a single QTL pair for each pair of chromosomes and assigned the QTL peak as the marker position with the smallest p value. We delineated the QTL region as the range of SNP markers with p-values within 25% of the peak in log-space: *p* < *p*_min_^(1 - 0.25)^.

Interacting QTLs were reported from the above scan only. We conducted a similar epistasis scan using R/qtl with standard two-locus interval mapping, the densest genetic map grid (*step* = 0), and ignoring same-chromosome comparisons (*clean*.*output* = *TRUE*; *clean*.*distance* = 999). One thousand permutations were used to obtain a null distribution for the interaction LOD score, and the 95^th^ percentile was used as the significance threshold. This scan found a total of 177 pairs of interacting QTLs for 21 novel isoforms; however, all of these QTL-pairs were suspect. For a large proportion of cases, it was clear that outliers had inflated statistical significance because the genotype group with the largest novel splicing proportion was represented by only one sample. This issue is likely specific to our dataset: with 10% of markers heterozygous in the mapping population, the two-locus genotypes that included heterozygotes (*AABḂ, AȦBB, ȦȦBḂ, AȦḂḂ*, and *AȦBḂ*) were usually represented by at most one sample. For the subset of QTL-pairs that did have sufficient sample sizes in each genotype group, the detected interaction appeared to be an artifact of missing data imputation (interval mapping involves the calculation of genotype probabilities for missing genotypes). Each of these QTLs had many genotypes missing; for some of the identified QTLs, all transgressive samples had missing data at both loci. We concluded that this method was inappropriate given the particular attributes of the dataset.

### Associations with individual spliceosome components

Sequences for *Arabidopsis thaliana* spliceosome protein and RNA components were obtained from KEGG and arabidopsis.org. Spliceosome sequences were aligned to our sunflower transcriptome using TBLASTN (McGinnis and Madden, 2004), and the best hit with e-value < 10^−20^ was retained for each component. The total number of tests was 4,275. With a Bonferroni correction, the significance threshold for this analysis was p < 1.16 × 10^−5^.

When a novel isoform was associated with both a spliceosome component and a regulatory QTL, we partitioned the variance explained using a linear model including all spliceosome components and QTLs associated with the novel isoform as predictors. For epistatic QTL-pairs, only the interaction term was included in the model. Interaction terms were included for each possible spliceosome component-QTL combination. We partitioned the sum of squares explained using eta-squared with type three sum of squares. To examine whether a QTL explains more variance in novel isoform splicing or expression of spliceosome components, we used individual linear models including one predictor and one response variable: the QTL was the predictor, and individual spliceosome components or the novel isoform phenotype was the response.

## Acknowledgments

This work utilized both the BioFrontiers Computing Core at the University of Colorado at Boulder (UCB) supported by BioFrontiers IT, and the RMACC Summit Supercomputer supported by the National Science Foundation (awards ACI-1532235 and ACI-1532236), UCB, and Colorado State University.

## Data availability statement

Sequence data reported in this paper are available as part of the Sequence Read Archive: Bioproject no. PRJNA417714.

## Author contributions

Conceptualization, original draft preparation, and review and editing were contributed by CCRS, LHR, BSH, and NCK. Formal analysis was performed by CCRS and NCK.

